# Modulation of fibroblasts phenotype by colorectal cancer cells-secreted factors is mostly independent of oncogenic KRAS

**DOI:** 10.1101/2022.07.05.498793

**Authors:** Patrícia Dias Carvalho, Susana Mendonça, Flávia Martins, Maria José Oliveira, Sérgia Velho

**Affiliations:** i3S - Instituto de Investigação e Inovação em Saúde, Universidade do Porto, Portugal; IPATIMUP – Instituto de Patologia e Imunologia Molecular da Universidade do Porto, Porto, Portugal; ICBAS - Institute of Biomedical Sciences Abel Salazar (ICBAS), University of Porto, Porto, Portugal; Department of Pathology, FMUP – Faculty of Medicine of the University of Porto, Porto, Portugal; INEB – Institute of Biomedical Engineering, University of Porto, Porto, Portugal

**Keywords:** Cancer-associated Fibroblasts, Colorectal cancer, Mutant KRAS, Secreted factors

## Abstract

KRAS mutations have been shown to extend their oncogenic effects beyond the cancer cell, influencing the tumor microenvironment components. Herein, we studied the impact of mutant KRAS on the modulation of cancer-associated fibroblasts (CAFs) pro-tumorigenic properties. To do so, we challenged CCD-18Co normal-like colon fibroblasts with control media (DMEM alone and DMEM+rhTGFβ1 media), and conditioned media from control and KRAS silenced colorectal cancer (CRC) cells. Two mutant KRAS CRC cell lines-HCT116 and LS174T, were used. Major pro-tumorigenic fibroblasts phenotypic features, such as α-SMA expression, TGFβ1 and HGF production, extracellular matrix components and metalloproteinases expression, collagen contraction and migration capacities, were analyzed upon fibroblast challenging with cells conditioned media. Our results showed that the mutant KRAS CRC cells-secreted factors are capable of turning normal-like fibroblasts into CAF-like by modulating α-SMA expression, TGFβ1 and HGF production and migration capacity, though in a cell line-specific manner. In this scenario, oncogenic KRAS showed to play a secondary role, regulating only discrete features in each cancer cell-educated fibroblasts. For instance, in HCT116, KRAS impairs fibroblasts migration and in LS174T it promotes α-SMA expression. In summary, our work suggests that mutant KRAS does not play a major role in controlling the CRC cell secreted factors that modulate fibroblasts behavior. This KRAS-independent modulation of fibroblast pro-tumorigenic features is likely to negatively impact the response to KRAS inhibitors, thus standing as a putative mechanism of resistance to KRAS-inhibition, with possible therapeutical relevance.

## INTRODUCTION

KRAS is the oncogene most frequently mutated in cancer, featuring on the group of key cancer targets. Notwithstanding mutations were discovered nearly 40 years ago, it remains one of the few that still lacks efficient therapies ^1^. Mutations occur in numerous cancer types at variable frequencies, prevailing in lung (32%), colorectal (CRC; 50%) and pancreatic (88%) cancers ^2^. They render KRAS constitutively active, thus fueling the cancer cell with strong mitogenic, survival, stemness, invasive, and pro-metastatic signals ^3,4^. Adding to this major oncogenic role, mutant KRAS (mutKRAS) is also a predictive factor of resistance to anti-EGFR targeted therapies ^5^.

Adding to the roll of key cancer activities controlled by mutKRAS, it has been shown that its oncogenic signaling extends beyond the cancer cell to orchestrate a pro-tumorigenic microenvironment ^6–11^. Within the tumor microenvironment (TME), immune cells and fibroblasts are amongst the most commomly cell types found ^6,12^. The effect of mutKRAS on the regulation of the tumor immune microenvironment is one of the most well described non-cancer cell effect. Specifically, mutKRAS immunomodulator effects, have been reported to occur at the level of recruitment, activation, and differentiation of immune cells, supporting the immune compartment protumorigenic roles, and cancer cell evasion from immunosurveillance ^6–11,13^. The effects of mutKRAS on the modulation of fibroblasts and cancer associated fibroblasts (CAFs) phenotypes are not so well studied and the majority of the available studies are focused on pancreatic cancer ^6,7^. Still, the existing data suggest that KRAS mutations in cancer cells can alter the behavior of fibroblasts, which in return affects both the cancer cells and the microenvironment through extracellular matrix (ECM) changes and growth factor signaling, contributing to tumor progression ^6,14^.

CAFs are an heterogeneous and plastic population, originating in their majority from the “activation” of local tissue fibroblasts, for instance by the action of tumor cells secreted factors, such as transforming growth factor-β (TGF-β) or platelet-derived growth factor (PDGF) ^15,16^. Together with the expression of some markers, such as α-smooth muscle actin (α-SMA) and fibroblast activation protein-α, among others, this activation state foresees the acquisition of contraction, proliferation, ECM synthesis/remodeling capacities, and a highly secretory phenotype ^15^. These characteristics make CAFs major modulators of the TME ^15,16^. CAFs participate in the acquisition and maintenance of most cancer hallmarks, such as sustaining proliferative signaling, deregulating cellular energetics, activating invasion an d metastasis and inducing angiogenesis being active players the construction of an immune permissive environment that supports cancer progression ^12,17^. In CRC, CAFs represent the most abundant stromal population ^12^. They are found at the tumor invasive front ^18,19^ and their abundance is a poor prognosis factor ^20^. Hence, given the central roles of CAFs in CRC progression and malignancy, studies approaching CRC cells-fibroblasts crosstalk, are still needed. Moreover, a better understanding of how mutKRAS affects the TME may reveal potential therapeutic targets that impair mutKRAS cancer cells by reverting the pro-tumorigenic properties of the TME components.

Herein, we aimed to uncover the effect of secreted factors from mutKRAS CRC cells on the modulation of fibroblasts properties. Our data revealed that fibroblasts properties were modulated by the cancer cells secreted factors, mostly independently of KRAS and in a cancer cell-dependent context. Therefore, our work sheds light on a potential mechanism of resistance to KRAS inhibition: upon KRAS inhibition, cancer cells continue to promote pro-tumorigenic properties on fibroblasts which on their turn may support the survival of cancer cells through KRAS-independent mechanisms.

## MATERIALS AND METHODS

### Cell culture

HCT116 CRC cell line and CCD-18Co normal colon fibroblasts were purchased from American Type Culture Collection. The CRC cell line LS174T was kindly provided by Dr. Ragnhild A. Lothe (Oslo University Hospital).

HCT116 was cultured in RPMI 1640 medium (Gibco, Thermo Fisher Scientific) and LS174T and CCD-18Co were cultured in DMEM medium (Gibco, Thermo Fisher Scientific). For all cell lines the respective medium was supplemented with 10% fetal bovine serum (Hyclone) and 1% penicillin–streptomycin (Gibco, Thermo Fisher Scientific). Cells were maintained at 37 °C in a humidified atmosphere with 5% CO2.

### Gene silencing with siRNA and conditioned medium production

HCT116 (150,000 cells/well) and LS174T (200,000 cells/well) were seeded in six-well plates and transfected after approximately 16h. Transfection was performed using Lipofectamine RNAiMAX (Invitrogen, Thermo Fisher Scientific) in reduced-serum Opti-MEM medium (Gibco, Thermo Fisher Scientific), following manufacturer recommendations. KRAS silencing was achieved using a specific ON-TARGETplus SMARTpool small interfering RNA (L-005069-00-0010; Dharmacon) at a final concentration of 10nM, and a non-targeting siRNA (D-001810-01-50; ON-TARGETplus Non-targeting siRNA #1; Dharmacon) was used as negative control. Forty-eight hours after transfection, cell monolayers were washed twice, and the respective serum-free culture medium was added. After additional 24h, the conditioned media (CM) from all the conditions were harvested, centrifuged to remove cell debris, filtered through a 0,2μm filter and stored at -20ºC until use. Total protein was extracted and KRAS silencing efficiency was evaluated by western blot (Supplementary Figure 1).

### CCD-18Co treatment with conditioned medium

CCD-18Co fibroblasts (100,000 cells/well) were seeded in six well plates. When approximately 90% confluence was reached (time 0), cells were washed twice and the CM from HCT116 and LS174T-control (siCTRL) and KRAS silenced (siKRAS) was added. At the same time, DMEM + 1% penicillin/streptomycin, and DMEM + 1% penicillin/streptomycin + 10 ng/mL rhTGFβ1 (Immunotools, GmbH) were used as negative and positive controls, respectively. After four days in culture, subsequent experiments were performed. Media from all conditions were collected, centrifuged to remove cell debris and stored at -20ºC. Total protein and mRNA were extracted from each condition.

### Protein extraction and Western blotting

Total protein was extracted using ice cold RIPA lysis buffer (50 mM Tris HCl; 150 mM NaCl; 2 mM EDTA; 1% IGEPAL CA-630; pH=7.5) supplemented with a protease inhibitor cocktail (Roche) and a phosphatase inhibitor cocktail (Sigma-Aldrich). Protein concentration was determined using the DCProtein assay kit (Bio-Rad). Briefly, 15μg of protein were resolved on sodium dodecyl sulphate-polyacrylamide gel electrophoresis (SDS-PAGE) under denaturing conditions and transferred to Protran Premium NC 0.45μm membranes (Amersham Biosciences, GE Healthcare). Membranes were blocked for 1h at room temperature (RT) and incubated overnight at 4ºC with agitation, with the respective primary antibody (KRAS-LS-Bio, LS-C175665; 1:4000; α-SMA-Abcam, ab7817, 1:250; GAPDH-Santa Cruz Biotechnology, sc-47724, 1:10000), diluted in 5% non-fat milk in PBS+ 0.5% Tween 20 (Sigma-Aldrich). Membranes were subsequently incubated for 1h at RT with anti-mouse (NA931, GE Healthcare) HRP-conjugated secondary antibody, and bands were detected using ECL (Bio-Rad) and autoradiography film (Amersham Biosciences, GE Healthcare) exposure. Band intensity was quantified using Fiji Software.

### mRNA expression analysis by qRT-PCR

Total RNA was extracted using the TripleXtractor (Grisp Research Solutions) following the manufacturer instructions. Briefly, cells were lysed in Triplextractor and after aqueous and organic phase separation using chloroform (MERCK), RNA was precipitated using isopropanol (Fisher Chemical). After 2 rounds of washes with 70% ethanol (Fisher Chemical), RNA was eluted in ultrapure RNases/DNases free water (Invitrogen, Thermo Fisher Scientific). A total of 0,5μg of RNA were converted to cDNA using qScript™ cDNA SuperMix, (Quantabio), according to the provided instructions. Quantitative-Real-Time-PCR (qRT-PCR) was performed in a 7500 Fast Real-Time PCR system (Applied Biosystems, Thermo Fisher Scientific) using TaqMan® Universal PCR Master Mix, No AmpErase® UNG (Applied Biosystems, Thermo Fisher Scientific) and standard TaqMan thermocycling conditions. TaqMan Gene Expression Assays (Thermo Fisher Scientific) or PrimeTime qPCR Assays (Integrated DNA Technologies) referred in supplementary Table 1 were used. Relative mRNA expression levels of the targeted genes were normalized to expression levels of the housekeeping gene GAPDH and estimated using the comparative 2(^−ΔCT^) method.

### Quantification of secreted factors by ELISA

The levels of total TGFβ1 (Legend Max, BioLegend) and HGF (RayBiotech) were quantified in the CM from HCT116 and LS174T cells (siCTRL and siKRAS) and in the media from all conditions collected at the final time-point of the experiment with CCD18-Co fibroblasts, following manufacturer instructions.

### Wound healing Assay

When CCD-18Co fibroblasts formed a confluent monolayer, a wound was made in using a 200μL tip. Detached cells were washed and controls and CM from HCT116 and LS174T siCTRL and siKRAS cells were added to the respective well. Two technical replicates/condition were performed. Wound healing was monitored every 24h until the final timepoint of four days. At all timepoints, two areas/wound were photographed using a Leica DMi1 inverted microscope with camera, using the 5x objective. Wound area was quantified using Fiji software.

### Collagen contraction Assay

After four days in culture, CCD-18Co fibroblasts were trypsinized, washed and a cell suspension of 3 × 10^5^ cells/mL was prepared in the appropriate control media or CM. From this cell suspension, 400μL were mixed with 200μL of a 3 mg/mL collagen in 0.1% acetic acid solution (rat tail type I; Merck Millipore), followed by the addition of 10μL of 1 M NaOH. As cell-free negative controls, 400μL of each medium were mixed with the collagen solution. From each mixture, 500μL were plated in a 24-well plate, allowing the gels to solidify at RT for 1h. After solidification, 600μL of the respective serum-free control media or CM were added and the gels were completely dissociated from the wells by gently running a pipet tip along gel edges. The plate was maintained in optimal culture conditions during 8h. Images of the plate were taken in a Bio-Rad Gel-Doc Imager and the final area of each gel was measured using Fiji Software.

### Statistical analysis

Statistical analyses were performed using the GraphPad Prism Software (version 9.0.0). Results are expressed as mean ± standard deviation (SD) and the specific performed test, as well as the number of independent biological replicates, are referred on each figure legend.

## RESULTS

### CRC cells secreted factors influence the expression of α-SMA and the secretion of TGFβ1 and HGF by CCD-18Co fibroblasts

In order to evaluate if oncogenic KRAS signaling extends to the outside of the cancer cells affecting the properties of fibroblasts, we evaluated the activation state of fibroblasts in response to cancer cell secreted factors. To do so, CCD-18Co fibroblasts were treated with CM from CRC cells-siCTRL and siKRAS, and control media (with and without rhTGFβ1). After four days of treatment, total protein was extracted and the expression of the fibroblast-activation marker α-SMA was evaluated. As expected, rhTGFβ1 treatment resulted in a significant increase of α-SMA protein expression. Similarly, CM from HCT116 cells was able to increase the expression levels of α-SMA, an effect that was independent of KRAS silencing. Even though to smaller levels, CM from LS174T siCTRL cells was also capable to increase the expression of this marker, whereas CM from LS174T siKRAS failed to reach statistical significance (Figure 1A and B). Attempting to explain these results, total TGFβ1 was quantified in the CM from CRC cells. The results showed that both HCT116 and LS174T cells secreted similar levels of TGFβ1 when comparing siCTRL with siKRAS conditions. Notably and in accordance with the levels of activation observed, TGFβ1 levels were higher in HCT116-derived CM in comparison to LS174T-derived CM (Figure 1C). Moreover, challenging of fibroblasts with CM, resulted in a significant increase of TGFβ1 levels in the media for the conditions with HCT116 siCTRL and siKRAS CM, while CM from LS174T cells showed no significative effect (Figure 1C).

**Figure 1.**
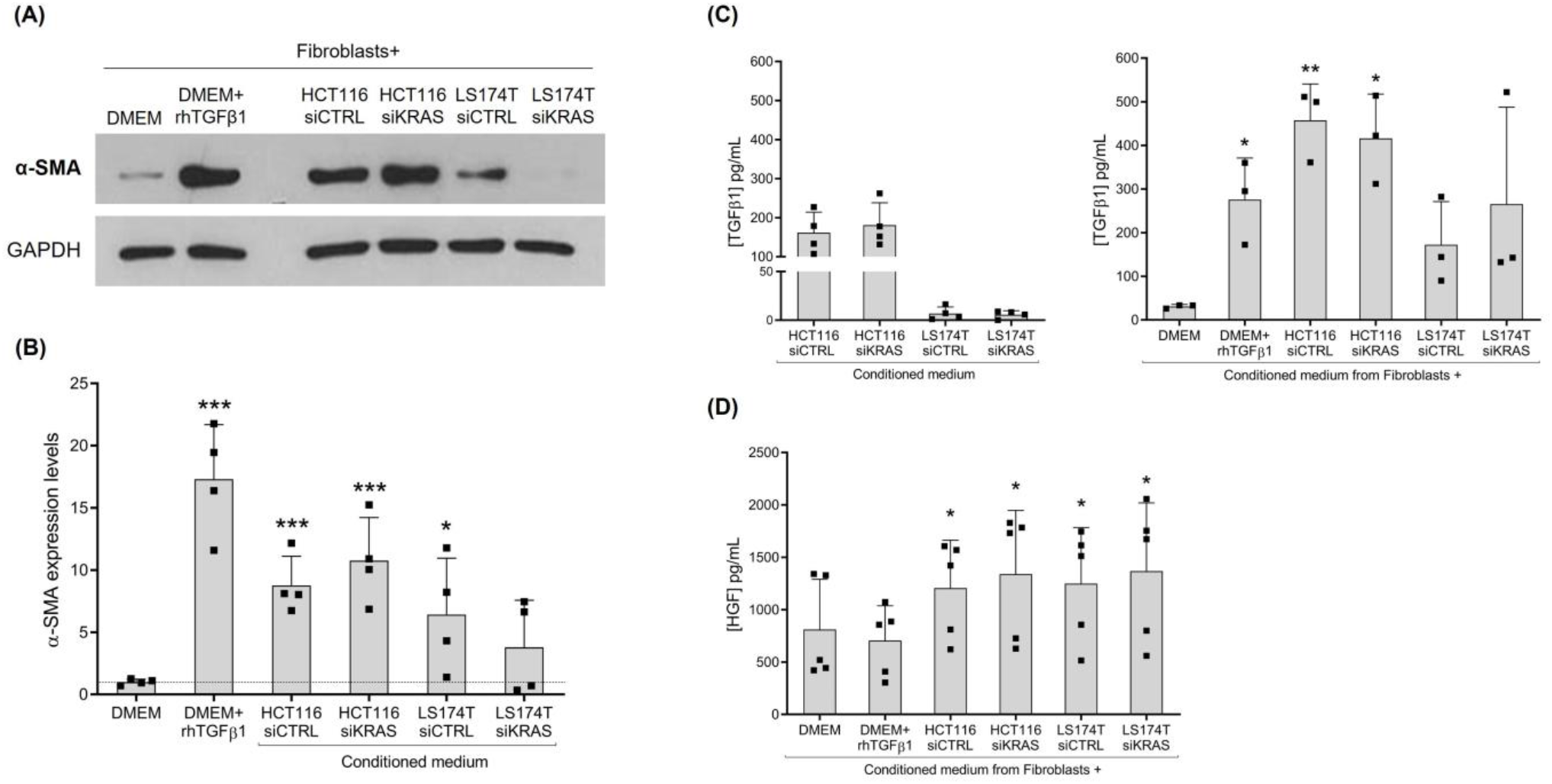
Culture with conditioned media (CM) from colorectal cancer cells is capable to influence the expression levels of α-SMA and production of TGFb1 and HGF by CCD-18Co fibroblasts. (A) Representative western blot illustrating α-SMA expression levels, and (B) the respective quantification of four independent biological replicates. (C) Levels of total TGFβ1 in the CM from HCT116 and LS174T siCTRL and siKRAS cells, as well as in the media collect from fibroblasts upon 4 four days of culture with control media (DMEM and DMEM+rhTGFβ) and HCT116 and LS174T (siCTRL and siKRAS) cells CM. (D) Levels of HGF produced by fibroblasts upon 4 four days of culture with control media (DMEM and DMEM+rhTGFb) and HCT116 and LS174T (siCTRL and siKRAS) cells CM. Values of the independent experiments performed are plotted as mean±SD. Data normality was tested using the Shapiro-Wilk test and a parametric or non-parametric t-test was performed in accordance, comparing all conditions with the DMEM control (*p≤0.05; **p≤0.01; ***p≤0.001).

Previous work from our group ^22^ and others ^23^ demonstrated that HGF is an important fibroblast-secreted pro-invasive factor. Also, HGF has been reported to induce the activation of fibroblasts, supporting their tumor promoting characteristics ^24^. Hence, the levels of this growth factor were also evaluated. HGF was not detected in the CM from CRC cells alone. In accordance with our previous results ^22^, non-activated and rhTGFβ1-activated fibroblasts produce similar levels of HGF. Though, challenging of fibroblasts with CM resulted in a significant increase in HGF production in all conditions (Figure 1D).

Together, this set of results demonstrates that the impact of CRC cells secretome on fibroblasts activation (α-SMA expression) and TGFβ1 and HGF secretion is mostly independent of oncogenic KRAS, except for LS174T in which α-SMA expression was dependent on KRAS.

### CRC cells secretome have no influence on the mRNA expression of extracellular matrix components but alters the expression of MMPs

Since the major role fibroblasts is the production and remodeling of ECM, the expression of some ECM components-fibronectin I and collagens type I, III and IV, and MMPs-1, 2, 3, 9 and 14, was evaluated in total mRNA from fibroblasts. Regarding the ECM components, qRT-PCR results show that activation with rhTGFβ1 results in a significative upregulation of *Fn1, Col1A1* and *Col4A1*, having no effect on the expression of *Col3A1*. None of the CRC cells-CM showed to influence the expression of the analyzed ECM components (Figure 2A). Concerning the expression of MMPs, rhTGFβ1 showed a tendency to increase the expression of *MMP2* and *MMP3*, having no impact on the expression of *MMP1* and *MMP14*. In opposition, CM from HCT116 cells-both siCTRL and siKRAS, downregulated the expression of *MMP 1, 2, 3* and *14*. LS174T siCTRL CM revealed to downregulate only *MMP2* and *14* and LS174T siKRAS CM only reached statistical significance for *MMP14* (Figure 2B). The expression of *MMP9* was not detected in none of the tested conditions.

**Figure 2.**
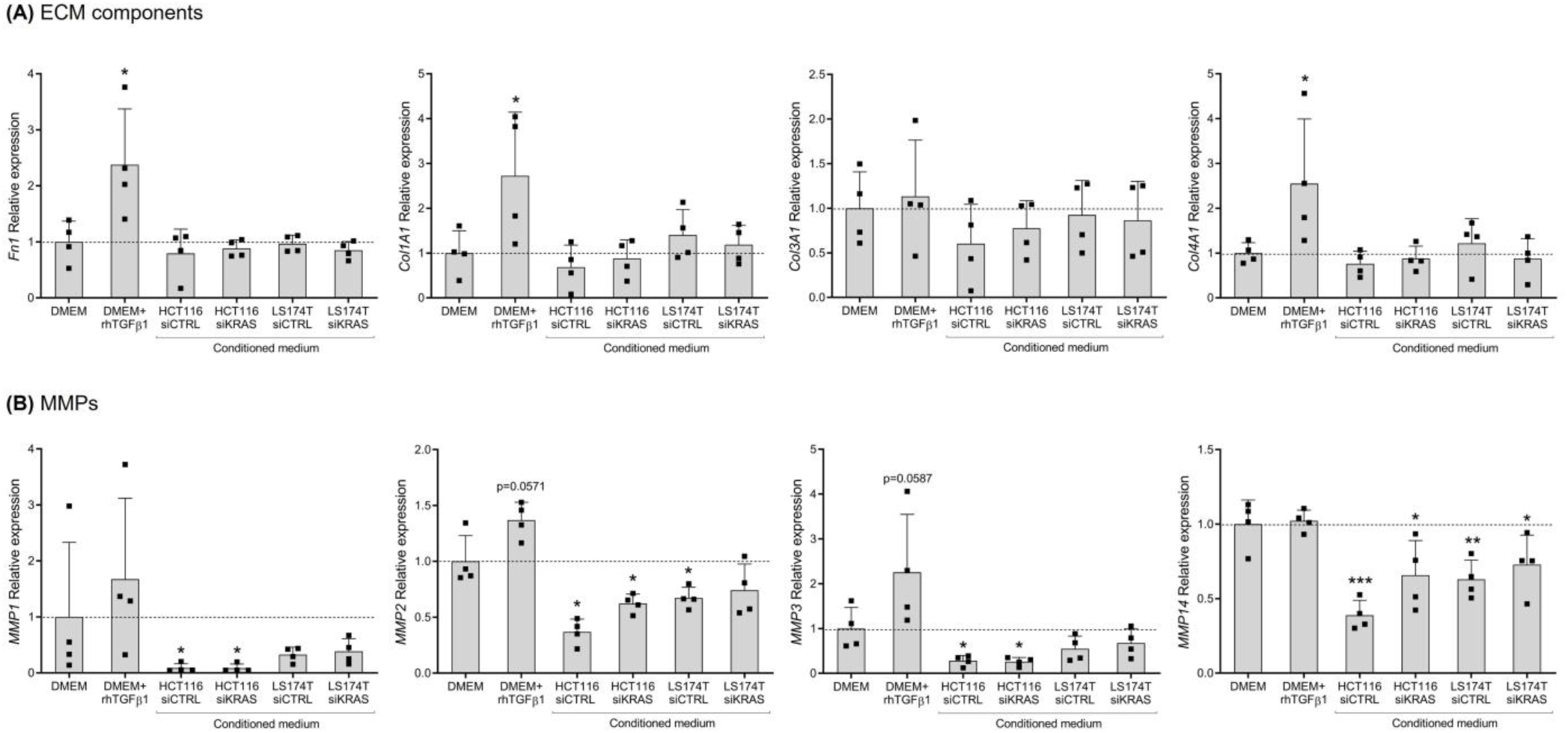
Conditioned media from CRC-cells have no impact on the expression of ECM components, while showing some effects on the expression of MMPs. (A) Relative expression levels of ECM components-*Fn1, Col1A1, Col3A1* and *Col4A1* and (B) *MMP*s-*1, 2, 3* and *14*. Values of the independent experiments performed were normalized to the DMEM control and are plotted as mean ±SD. Data normality was tested using the Shapiro-Wilk test and a parametric or non-parametric t-test was performed in accordance, comparing all conditions with the DMEM control (*p≤0.05; **p≤0.01; ***p≤0.001).

This data show that CRC cells secreted factors do not influence the mRNA levels of the analyzed ECM components. However, all the detected MMPs, showed to be downregulated by HCT116 cells-secreted factors, independently of KRAS. LS174T cells-CM only downregulated the expression of *MMP2* and *14*, though only *MMP2* downregulation showed and association with the presence of KRAS.

### HCT116 and LS174T secreted factors differentially affect fibroblast collagen-contraction and migration capacities

Since contraction and migratory capacities are also important features of activated-fibroblasts, we evaluated these capacities. Contraction capacity was evaluated by assessing the changes in the area of fibroblast-populated collagen type I gels, upon “education” of fibroblasts with the respective control media and CRC cells-CM for four days. Collagen gels with rhTGFβ1 activated-fibroblasts showed a decreased area when compared to the control (DMEM alone), demonstrating that activation with rhTGFβ1 increases the contraction capacity of CCD-18Co fibroblasts. CM from HCT116 cells-siCTRL and siKRAS, had no effect on collagen area when compared with the DMEM control. Contrarywise, collagen gels with CM from LS174T siCTRL and siKRAS cells showed an increased area when compared to the DMEM control, indicating a decreased fibroblast contraction capacity (Figure 3 A and B).

**Figure 3.**
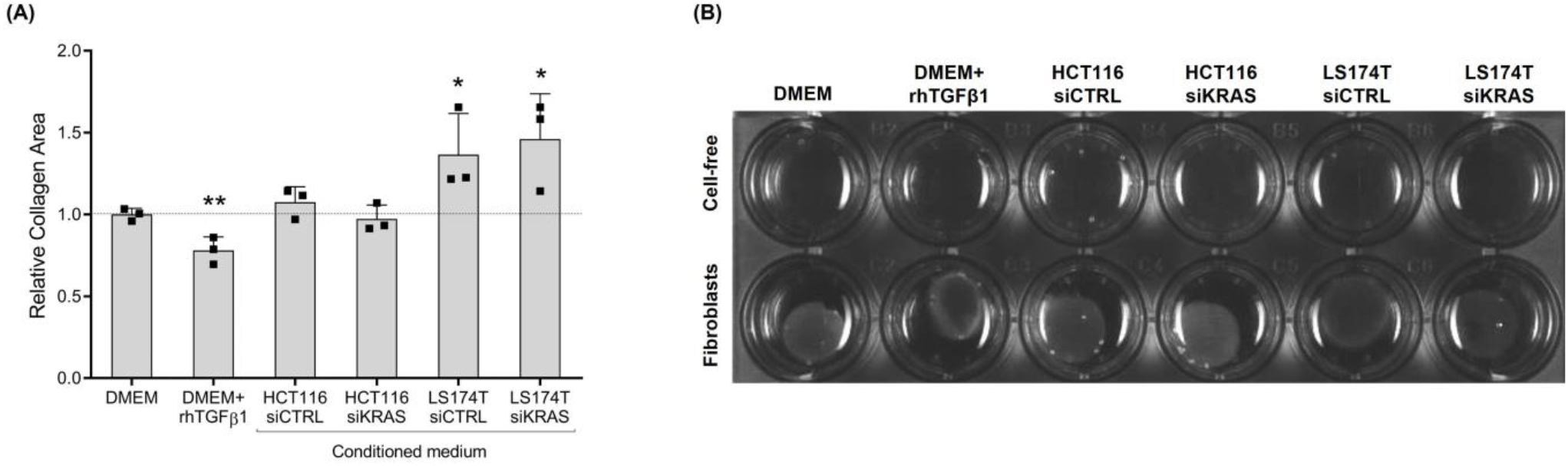
rhTGFβ and conditioned medium from LS174T cells are capable to influence fibroblast-contraction capacity. (A) After 4 days in culture with control media (DMEM and DMEM+rhTGFβ1) and HCT116 and LS174T (siCTRL and siKRAS) cells CM, pre-educated fibroblasts were embedded in collagen type I and relative area of the collagen pads was quantified after 8h. Collagen gels with rhTGFβ1 activated-fibroblasts presented a decreased area, demonstrating fibroblast contraction capacity. CM from HCT116 cells-siCTRL and siKRAS, had no effect on collagen area. Collagen gels with CM from LS174T siCTRL and siKRAS cells showed an increased area, indicating a decreased fibroblast contraction capacity. Values of the independent experiments performed were normalized to the DMEM control and are plotted as mean ±SD. Data normality was tested using the Shapiro-Wilk test and a parametric or non-parametric t-test was performed in accordance, comparing all conditions with the DMEM control (*p≤0.05; **p≤0.01; ***p≤0.001). (B) Representative image showing cell-free gels, lacking contraction and contracted fibroblast-populated gels.

To evaluate the impact of CRC cells-secretome on migration capacity, we resorted to a wound-healing assay, in which we evaluated the wound-area during the course of the four days. Surprisingly, the results showed no differences in the migration of fibroblasts cultured in DMEM alone or in DMEM+rhTGFβ1. However, CM from HCT116 siKRAS cells showed to significantly increase the migration capacity, when compared to DMEM alone in all time-points and from time-point 48h forward when compared to DMEM+rhTGFβ1. Furthermore, this effect was verified when comparing to the migration rate induced by CM from HCT116 siCTRL cells-CM (from time-point 48h forward), demonstrating a KRAS dependency. On the contrary, independently of KRAS, fibroblasts in LS174T cells-CM, also migrate faster than DMEM and DMEM+rhTGFβ1 (siCTRL at 72h and 96h; siKRAS from 48h forward) (Figure 4 A and B).

**Figure 4.**
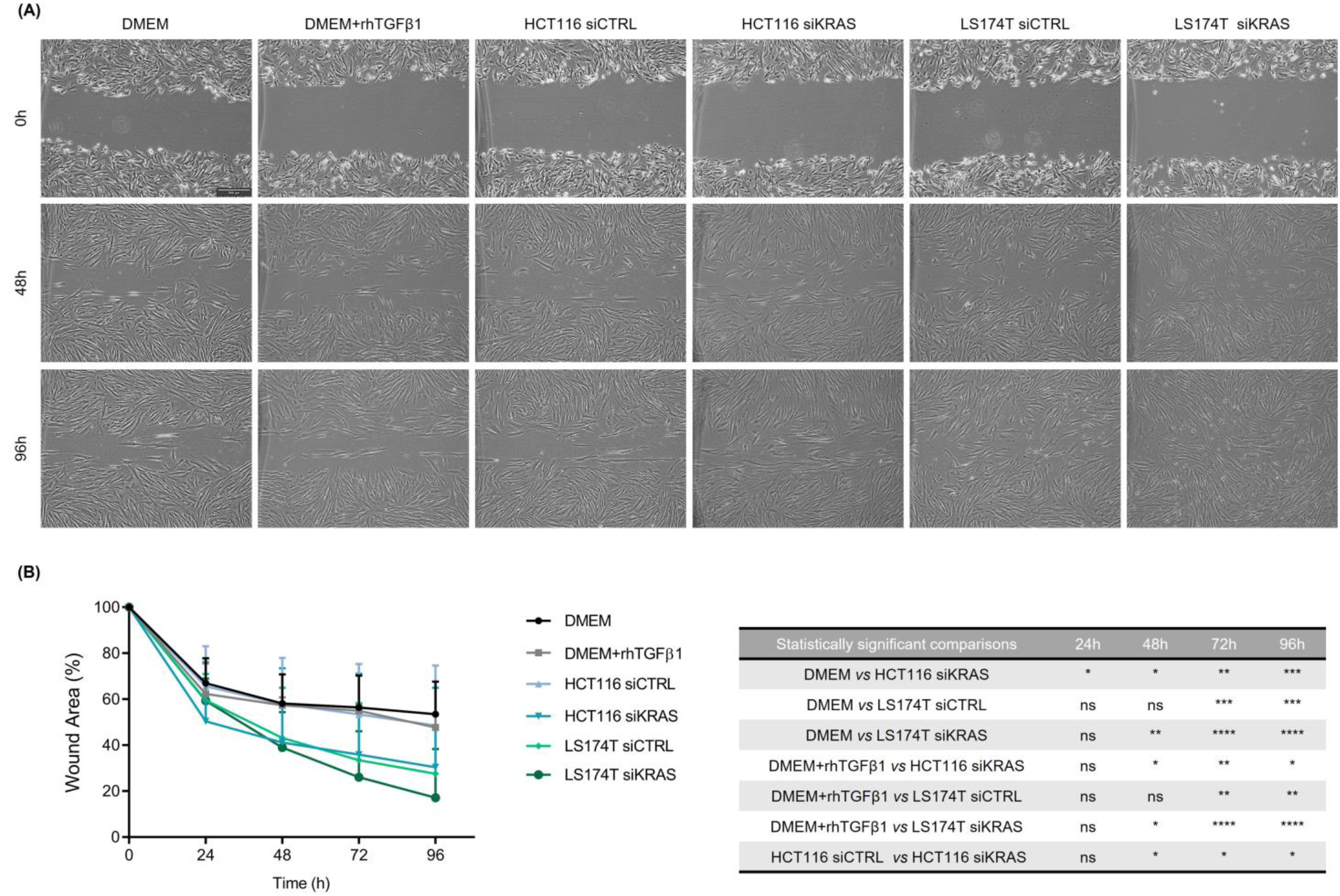
Conditioned media from HCT116 siKRAS and LS174T siCTRL and siKRAS increased the migration capacity of fibroblasts. When a confluent monolayer was formed, a wound was made and control media (DMEM and DMEM+rhTGFb) and HCT116 and LS174T (siCTRL and siKRAS) cells CM were added to the respective condition. Wounds were monitored every 24h during four days. (A) Representative micrographs of all conditions at time-points 0h, 48h and 96h. All images were acquired using the 5x objective, scale bar corresponds to 500μm. (B) Quantification of the wound area along time and the respective statistically significant differences. No differences were found in the migration of fibroblasts cultured in DMEM alone or in DMEM+rhTGFβ1. CM from HCT116 siKRAS cells significantly increased fibroblast migration capacity, when compared to DMEM alone in all time-points and from time-point 48h forward when compared to DMEM+rhTGFβ1 and HCT116 siCTRL cells-CM. LS174T cells-CM (siCTRL and siKRAS) increased fibroblast migration capacity when compared to DMEM and DMEM+rhTGFβ1 (siCTRL at 72h and 96h; siKRAS from 48h forward). Three independent biological replicates were performed. Values are plotted as mean ±SD of 3 biological replicates and statistical significance was evaluated using the two-way ANOVA considering repeated measures by both factors, with Tukey’s multi comparison test (*p≤0.05; **p≤0.01; ***p≤0.001; ****p≤0.0001; ns: not significant).

In sum, these results show that CM from HCT116 cells had no impact on the contraction capacity of fibroblasts, however CM from KRAS silenced cells was able to increase fibroblasts-migratory capacity. On their turn, CM from LS174T cells decreased fibroblasts-contraction capacity while increasing their migratory ability, being both effects independent of KRAS.

## DISCUSSION

Cancer cells and fibroblasts within the TME are known to establish a crosstalk that is essential for tumor progression and malignancy. One of the routes involved in this interaction occurs *via* secreted factors by both cell types that are able to modulate each other properties ^15,16,25^. Though, the impact of cancer cell-specific alterations, namely KRAS oncogenic activation, on the regulation of this crosstalk is poorly explored. We have recently reported that aproximatly 2/3 of mutant KRAS-associated proteome is regulated by fibroblasts-secreted factors ^21^, impacting processes such as cell invasion ^22^. Herein, we addressed how mutKRAS impacts fibroblast properties. Our data revealed that, in contrast to what has been described regarding the capacity of mutKRAS cancer cells to modulate the tumor immune microenvironment, the pro-tumorigenic features of fibroblasts are mainly regulated by CRC cells independently of oncogenic KRAS.

In analogy to tissue myofibroblasts, major features of the “activated” state, rely on the expression of activation markers, acquisition of contraction and migration capacities and a secretory phenotype involving the production of growth factors and cytokines, ECM components and remodeling enzymes ^15,16^. Among the factors that can activate fibroblasts, TGFβ is recognized as a major inducer ^26–28^. TGFβ signaling is related to the expression of the cytoskeleton protein α-SMA, one of the most widely accepted fibroblast-activation markers, as well as with ECM construction and remodeling activities ^16,28–30^. Additionally, in CRC, enhanced TGFβ signaling has been identified and associated with a stromal signature that correlates with poor prognosis ^31^ and stromal TGFβ has been shown to be essential for metastasis formation ^32^. In this work, activation of fibroblasts with rhTGFβ1 resulted in increased α-SMA protein expression, upregulation of *Fn1, Col1A1* and *Col1A4* and, even though not reaching statistical significance, there was a tendency to upregulate *MMP2* and *3*. More so, it was also capable to increase collagen contraction capacity, but failed to show an effect on the migration rate. In line with this results, HCT116 cells were shown to produce high levels of TGFβ1, similar between siCTRL and siKRAS cells, and the CM from these cells was in fact capable to increase the protein levels of α-SMA, as well as to contribute to the increased levels of TGFβ1 detected upon stimulation of fibroblasts with this CM, suggesting the existence of an autocrine positive feedback loop. The production of high levels of TGFβ1 by this cell line, with consequent increase of fibroblasts production of this growth factor and increased α-SMA expression, have also been reported by other authors ^30^. Despite the lower levels of TGFβ1 found, CM from LS174T siCTRL was also able to increase the levels of α-SMA while the CM from the siKRAS, presenting similar levels of TGFβ1, did not, showing that this may not be the only factor contributing to the increased expression of the activation marker analyzed. In accordance with the difference in TGFβ1 production between the two cell lines, is the fact that the cell line HCT116 belongs to the poor prognostic mesenchymal/stromal type CMS4, that has been characterized by upregulation of *TGFβ1/2* ^33^. Moreover, not all the subsequent observed effects can be correlated with the α-SMA expression levels, nor with the levels of secreted TGFβ1.

HGF is a well-known fibroblast-secreted pro-invasive factor in various solid cancer models ^23,34,35^ and our previous work showed that it can drive HCT116 and LS174T cells invasion, in a mutKRAS-dependent way ^22^. In addition to its pro-invasive role, stromal derived HGF has been reported to promote adhesion of CRC cells to endothelial cells, facilitating metastasis ^36^; to underlie resistance mechanisms to RAF inhibitors in a melanoma model ^37^ and to EGFR-targeting in different cancer types ^38,39^. Moreover, it has also been reported to induce the expression of fibroblast-activation markers in breast ^24^ and gastric cancer ^40^ models. Herein, the HGF quantification results showed that despite not producing this growth factor, HCT116 and LS174T secreted factors prompted fibroblasts to increase HGF secretion, an effect that was independent of KRAS. Our data further demonstrates that HGF production is independent of TGFβ1. The identification of the cancer cell-derived factors triggering HGF production by fibroblasts deserve further attention. TGFβ1-targeting therapies are being developed in an attempt to interfere with CAFs activation and signaling ^41^. However, according to our data, it is likely that TGFβ1 targeting will not be sufficient to abolish HGF production that, as already mentioned, is essential to drive cancer cell invasion.

Moreover, ECM remodeling is a major fibroblast-instructed activity impacting cancer cell invasion ^42,43^. TGFβ-activated fibroblasts increase their production of fibronectin and collagens as well as of MMPs, thus impacting ECM deposition and degradation, respectively ^28,30^. Our results showed that rhTGFβ1 was sufficient to upregulate *Fn1, Col1A1* and *Col4A1 expression* on CCD-18Co fibroblasts. However, CM from CRC cells, particularly from HCT116 which presented high levels of TGFβ1, did not affect the expression of the ECM components analyzed. Of note, using *in vitro* and *in vivo* models of rectal cancer, mutKRAS was recently shown to downregulate the expression of ECM components, such as *Fn1*, in fibroblasts. However, this effect was shown to require epithelial cells-fibroblasts direct co-cultures ^14^, a condition that was not performed in this work. Still, our data reinforce these findings by showing that mutKRAS have no impact on CRC cells-secreted factors that modulate the production of ECM components by fibroblasts. The same observation is likely to extend to the regulation of *MMPs* expression. TGFβ1 is reported to upregulate *MMPs* expression in CRC tissue-isolated fibroblasts (particularly, *MMP2, 3, 7, 9, 14, 15, 16* and *28*) ^30^. In addition, incubation of CRC tissue-isolated fibroblasts with CM from HCT116 cells was previously shown to increase fibroblasts expression of *MMP2, 3* and *14* ^30^. In contrast with these observations, in our work, stimulation of CCD18-Co naïve fibroblasts with rhTGFβ1 failed to significantly upregulate the expression of the *MMPs*, though a tendency to upregulate *MMP2* and *3* was observed. Contrarywise, CM from CRC cells (HCT116 siCTRL/siKRAS and LS174T siCTRL) lead to a downregulation of *MMPs (MMP1, 2 and 3 in HCT116 and MMP2 and 14 in LS174T)* expression in CCD-18Co fibroblasts. The different origins and state of activation of the fibroblasts used in our work and in the work of Hawinkels and colleagues^30^ may explain the discrepancy on the results. While we started from normal-like fibroblasts which were then stimulated with the different CRC cells CM, the previous work started from cancer-isolated fibroblasts. Additionally, of all the MMPs analyzed, we could not detect *MMP9* expression in none of the conditions. According to the literature, *MMP9* expression seems to require direct co-culture of fibroblasts with cancer cells ^44^, and a 3D culture system ^45^ in opposition to monolayer cultures as the ones used in our study.

TGFβ1 (along with increased α-SMA expression) is also reported to increase fibroblast-collagen contraction capacity in different models ^29,46^ and an increased α-SMA expression demonstrated to be sufficient to enhance different fibroblast models contractile activity ^47^. Our results regarding collagen contractility of fibroblasts stimulated with rhTGFβ1 corroborate these observations. However, a different scenario was found when fibroblasts were treated with cancer cells CM. Specifically, fibroblasts treated with CM from LS174T cells, presented slightly higher levels of TGFβ1 when compared to DMEM-only treated fibroblasts, though a decrease in fibroblast-contraction capacity was observed. Also, no changes on collagen contractility were observed when fibroblasts were stimulated with HCT116 (siCTRL and siKRAS) CM, despite the observed increase of TGFβ1 levels and significant upregulation of α-SMA expression relatively to DMEM-treated fibroblasts. The migratory capacity of fibroblasts was also differentially regulated among the different conditions tested. For instance, DMEM alone, DMEM+rhTGFβ1 and CM from HCT116 siCTRL cells did not promote any changes on fibroblast-migration capacity. Notably, CM from HCT116 KRAS-silenced cells caused a significant increase in the migratory capacity of fibroblasts, demonstrating that mutKRAS in HCT116 negatively regulates fibroblasts migration. This KRAS-dependent effect was cell line specific, as CM from LS174T siCTRL and siKRAS were equally capable to increase fibroblast-migration. Interestingly, other authors have shown that CM from KRASV12 transformed mouse colon cells increased the migration capacity of fibroblasts through the action of heparin-binding epidermal growth factor-like growth factor (HB-EGF), despite having no effect on α-SMA expression ^48^. So, these results raise the question on whether some of KRAS-orchestrated effects on fibroblasts are solely cell line-dependent or might be mutation-specific.

Overall, the discrepancies between TGFβ1-modulated and cancer cell CM-modulated fibroblasts properties found throughout our study highlight the potential that other factors present in the CM of the cancer cell lines have to skew fibroblasts properties towards different states or subpopulations. For instance, in pancreatic cancer, tumor secreted TGFβ and IL1 have been shown to underly the formation of two distinct fibroblasts subpopulations-the myofibroblastic CAFs and the inflammatory CAFs, respectively^49^. Likewise, as each cancer cell line induced distinct alterations on fibroblasts features (Figure 5), the study of the composition of the CM from these cell lines (growth factors, cytokines and extracellular vesicles) could help to gain further insights on the mechanisms governing fibroblasts heterogeneity within and between tumors.

**Figure 5.**
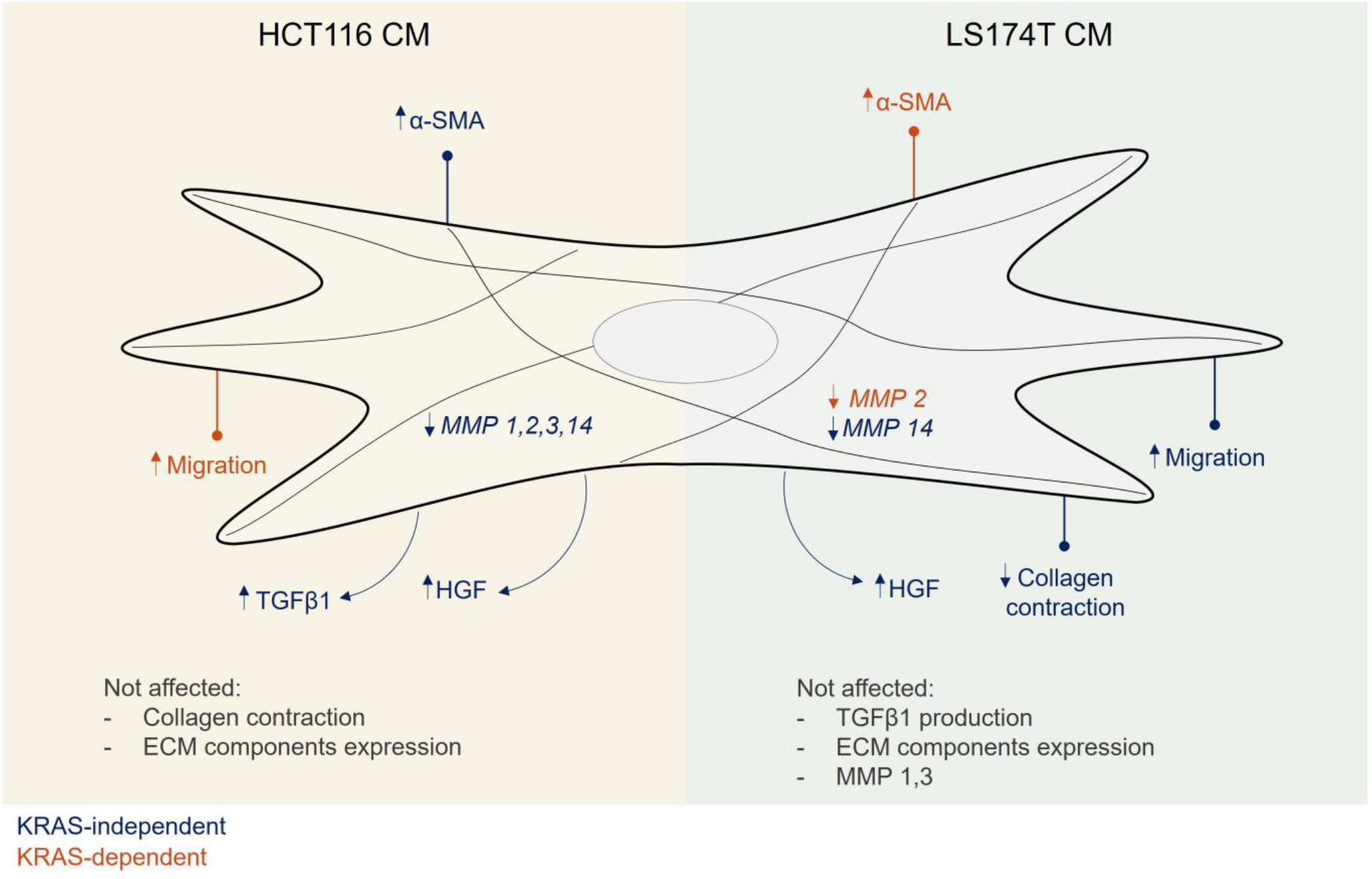
Conditioned media form HCT116 and LS174T cells modulate different aspects of fibroblast phenotype, mostly independently of KRAS. Secreted factors from HCT116 cells are able to increase fibroblast α-SMA expression and TGFβ1 secretion, as well as downregulate *MMP*s expression. Secreted factors from LS174T cells led to an increase of fibroblast migration and a decrease of collagen contraction capacities, and a downregulation of *MMP14* expression. Independently of KRAS, secreted factors from both cell lines increased the levels of fibroblast-secreted HGF. A KRAS dependency was only found regarding the capacity to regulate migration, in the case of HCT116 cells, and α-SMA and *MMP2* expression, in the case of LS174T cells.

In summary, our study demonstrates that mutKRAS does not play a major and general regulatory role on the secreted factors that modulate the analyzed fibroblasts properties. More so, each cancer cell line was able to modulate most of the fibroblasts’ features analyzed, in a cell line dependent manner. In this scenario, mutKRAS played a secondary role by regulating discrete features in each cancer cell-educated fibroblasts population. These observations raise a clinically relevant point regarding the response to the recently approved KRAS-targeted therapy and the search for combinatorial treatments to improve the response to KRAS inhibition: cells that survive KRAS inhibition are still able to promote pro-tumorigenic features on fibroblasts. In fact, the capacity revealed by KRAS inhibited CRC cancer cells to induce fibroblasts migration, activation, secretion of the pro-invasive factor HGF and the immunosuppressive and epithelial-to-mesenchymal inducer TGFβ1 is noteworthy. Therefore, there is a strong possibility that KRAS-inhibited cancer cell-educated CAFs support the high degree of tolerance to KRAS inhibition observed in tumors. More so, our study opens a new window of opportunity to improve responses to KRAS inhibition by understanding whether and how the cancer cell-educated fibroblasts contribute to support tolerance and resistance to KRAS inhibition and how we can therapeutically explore this potential vulnerability.

## Acknowledgments

The authors thank IPATIMUP’s Cell Lines Bank for cell line authentication.

## Author Contributions

Conceptualization: PDC, SV; Formal analysis: PDC; Funding acquisition: SV; Investigation: PDC, SM, FM; Methodology: PDC, SM, FM; Project administration: SV; Resources: MJO and SV; Supervision: MJO and SV; Validation: PDC, SM, FM; Visualization: PDC; Writing – original draft: PDC; Writing – review & editing: SM, FM, MJO, SV. All authors have read and approved the manuscript. The work reported in the paper has been performed by the authors, unless clearly specified in the text.

## Data Availability Statement

The data that support the findings of this study are available from the corresponding author upon reasonable request.

