## Supplementary Material for "Modulation of fibroblasts phenotype by colorectal cancer cells-secreted factors is mostly independent of oncogenic KRAS"

**Supplementary Table 1. qRT-PCR probes used in this work**

| Gene | Reference | Manufacturer |
|---|---|---|
| ***MMP1*** | Hs00899658_m1 | Thermo Fisher Scientific |
| ***MMP2*** | Hs01548727_m1 | Thermo Fisher Scientific |
| ***MMP3*** | Hs.PT.58.14758187 | Integrated DNA Technologies |
| ***MMP9*** | Hs00957562_m1 | Thermo Fisher Scientific |
| ***MMP14*** | Hs00237119_m1 | Thermo Fisher Scientific |
| ***FN1*** | Hs.PT.58.40005963 | Integrated DNA Technologies |
| ***Col1A1*** | Hs.PT.58.15517795 | Integrated DNA Technologies |
| ***Col3A1*** | Hs.PT.58.40254063 | Integrated DNA Technologies |
| ***Col4A1*** | Hs.PT.58.15679435 | Integrated DNA Technologies |
| ***GAPDH*** | Hs.PT.39a.22214836 | Integrated DNA Technologies |

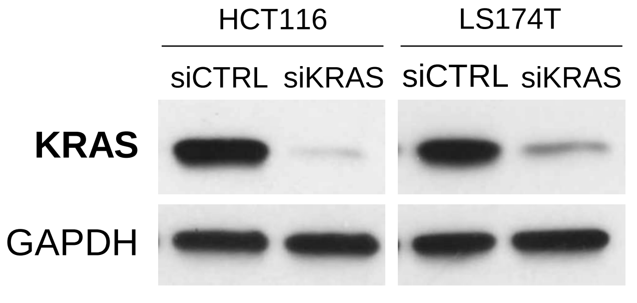


**Supplementary Figure S1. Representative Western Blots showing efficient KRAS silencing.**
